# Prioritizing ecological connectivity among protected areas in Colombia using a functional approach for birds

**DOI:** 10.1101/2023.09.12.557382

**Authors:** Daniela Linero Triana, Camilo Andrés Correa-Ayram, Jorge Velásquez-Tibatá

**Affiliations:** Audubon Americas, National Audubon Society, Bogotá, Colombia; Pontificia Universidad Javeriana, Bogotá, Colombia. Facultad de Estudios Ambientales y Rurales. Departamento de Ecología y Territorio

**Keywords:** Biodiversity conservation, connectivity network, multispecies connectivity, wildlife corridors, ecological restoration, landscape connectivity

## Abstract

Ecological connectivity among protected areas (PAs) is essential to improve biodiversity conservation and management effectiveness in the long term under global change. Developing strategic plans and identifying spatial priorities are practical actions for establishing and strengthening interconnected networks of PAs. In Colombia, this planning is fundamental to conserve its extraordinary bird diversity in the face of multiple threats, including climate change and deforestation. We develop a connectivity model focused on multiple bird species to identify critical sites to preserve the ecological connections among PAs in Colombia. Based on land cover data and expert knowledge, we created movement resistance surfaces and modeled Least-Cost Corridors among terrestrial PAs for 26 forest-dependent species. We also used circuit and least-cost models to locate conservation priorities and restoration opportunities, estimating the potential gain in connectivity with the Equivalent Connected Area (ECA) index. We found 581,531 km^2^ belonging to corridors among PAs for all focal species. Priority sites for movement within corridors covered 212,551 km^2^ and were predominantly located across Andean and Amazonian forests. Restoration opportunities covered 79,203 km^2^ and were concentrated in agricultural lands of the Andes and Caribbean regions. Restoring these areas could increase the national forest cover by 7% and connectivity by 14%. Our results provide a national-level assessment of functional connectivity priorities to maintain and improve the interconnections among PAs. This assessment could guide efforts related to conservation, restoration, and implementation of management tools that facilitate the movement of native species across transformed lands. These actions are crucial to meet the targets outlined in the post-2020 global biodiversity framework to achieve well-connected systems of PAs during this decade and until 2050.

## Introduction

Protected areas (PAs) are the primary strategy for safeguarding biodiversity worldwide and play a key role in sustaining ecological functions in the long term (Gaston et al., 2008; Pimm et al., 2018; Watson et al., 2014). However, the ability of individual PAs to protect species is limited because their sizes may be too small to support viable populations, and they are becoming increasingly isolated in landscapes dominated by anthropogenic lands (Chundawat et al., 2016; Defries et al., 2005; Naughton-Treves et al., 2005; Williams et al., 2022). Therefore, maintaining and restoring connectivity in PA networks is a priority to reverse biodiversity loss (Brennan et al., 2022; Hilty et al., 2020; Pulsford et al., 2015). Natural connections between PAs allow numerous biological processes, such as dispersal, migration, and gene flow, and are essential for species to respond and adapt to climate change (Heller & Zavaleta, 2009; Hilty et al., 2006, 2020; Schmidt et al., 2020; Schmitz et al., 2015). For these reasons, international policies such as the Convention on Biological Diversity (CBD; Secretariat of the Convention on Biological Diversity, 2020) and the post-2020 global biodiversity framework, emphasize the importance of enhancing connectivity among PAs. The framework explicitly calls for urgent action to establish well-connected PA systems during this decade (Convention on Biological Diversity, 2021).

Optimizing the connectivity of PA networks can be achieved through numerous planning strategies, including protecting and restoring corridors that facilitate species movements between habitat patches (Dutta et al., 2018; Heller & Zavaleta, 2009; Hilty et al., 2006). The most common approaches to optimize corridor placement are least-cost analysis and circuit theory (Adriaensen et al., 2003; Anantharaman et al., 2019; Correa Ayram et al., 2015; McRae et al., 2008). Both assume species incur costs when they move due to energy expenditures and mortality risks associated with different landscape features (Zeller et al., 2012). These costs vary between species depending on their ecological requirements and movement abilities and are usually expressed through resistance surfaces that describe the difficulty of crossing each grid cell in a landscape (Adriaensen et al., 2003; McClure et al., 2016; Zeller et al., 2012). Least-cost models use resistance surfaces to estimate the shortest and less costly corridors between patches assuming organisms have perfect knowledge of their surroundings (Adriaensen et al., 2003; McClure et al., 2016). Circuit theory complements this approach by identifying all possible paths between habitat patches that minimize movement resistance, assuming individuals behave as random walkers (McRae et al., 2008).

Using these approaches to identify corridors is crucial for Colombia, where PAs are a central strategy for halting the loss of the country’s exceptional biodiversity (Bonilla-Mejía & Higuera-Mendieta, 2019). Approximately 1,443 national, regional, and local PAs have been established, representing ca. 16% (190,900 km^2^) of the country’s terrestrial area (Parques Nacionales Naturales de Colombia, 2022). However, due to the accelerated land-use transformation in Colombia, PAs are frequently embedded in transformed lands, which may hinder movement across the landscape matrix for several species (Castillo et al., 2020; Murillo-Sandoval et al., 2022). Research has shown that the conversion of natural ecosystems in Colombia is significantly decreasing habitat connectivity in the Andean ecoregions, the Andes Amazon Transition Belt, and the national PA system (Castillo et al., 2020; Godínez-Gómez et al., 2021; Murillo-Sandoval et al., 2022). Furthermore, it is estimated that only 4.8% of Colombia’s land area is covered by connected national PAs (Godínez-Gómez et al., 2021). The low connectivity of PAs may increase the extinction probability of various populations and compromise the ecosystems’ flows and functioning (Haddad et al., 2023; Hansen & DeFries, 2007; Pulsford et al., 2015).

Consequently, protecting and improving connectivity between PAs in Colombia is a valuable action to support biodiversity conservation. In particular, Colombia is home to the largest avifauna in the world, with nearly 2,000 species presenting distinct habitat requirements and dispersal abilities (Echeverry-Galvis et al., 2022). Ensuring the movement of birds across the country is crucial for conserving this extraordinary diversity, as it may mitigate the adverse effects of forest fragmentation on altitudinal bird migration and the anticipated reduction and fragmentation of geographic ranges due to climate change (Hannah, 2008; Hsiung et al., 2018; Powell & Bjork, 1994; Velásquez-Tibatá et al., 2013).

Here, we developed a multispecies bird connectivity model to identify the places across the landscape matrix that may be critical to conserving and restoring connectivity among Colombian PAs. We selected 26 focal bird species that use forest habitats but exhibit various home range sizes and dispersal abilities. We modeled Least-Cost Corridors for each species based on expert criteria, species distribution models, and climate and land cover data. We then applied circuit theory models to identify pinch points or movement bottlenecks as priority conservation areas within the corridors. We also employed a least-cost approach to locate forest restoration opportunities with the potential to improve connectivity among PAs. We used individual species results to create multispecies composite maps highlighting spatial agreements for corridors, priority conservation areas, and restoration opportunities. Finally, we estimated the potential connectivity gain from forest restoration with the Equivalent Connected Area (ECA) index.

## Methods

### Study area

Colombia is in the northwestern end of South America (continental territory of ca. 1.1 million km^2^). As a result of the wide variety of topography, soils, and climate, Colombia is home to an immense diversity of species and ecosystems (Etter, 1993; Sánchez-Cuervo et al., 2012). Natural forests occupy around 52% of the national surface, extending from humid forests in the Amazon to high Andean forests (Etter, Andrade, Saavedra-Ramírez, et al., 2017; Instituto de Hidrología Meteorología y Estudios Ambientales - IDEAM, 2022). Historically, these ecosystems have been deforested mainly due to agricultural expansion, land grabbing, illicit crops, timber extraction, mining, and forest fires (Armenteras et al., 2013; Dávalos et al., 2016). The primary strategy to protect forests against these threats has been the National System of Protected Areas (SINAP; Figure 1). This system includes 1,443 terrestrial PAs under 15 different categories ranging from strict protection (e.g., National Parks) to sites under sustainable management (e.g., Civil Society Reserves; Parques Nacionales Naturales de Colombia, 2022).

**Figure 1.**
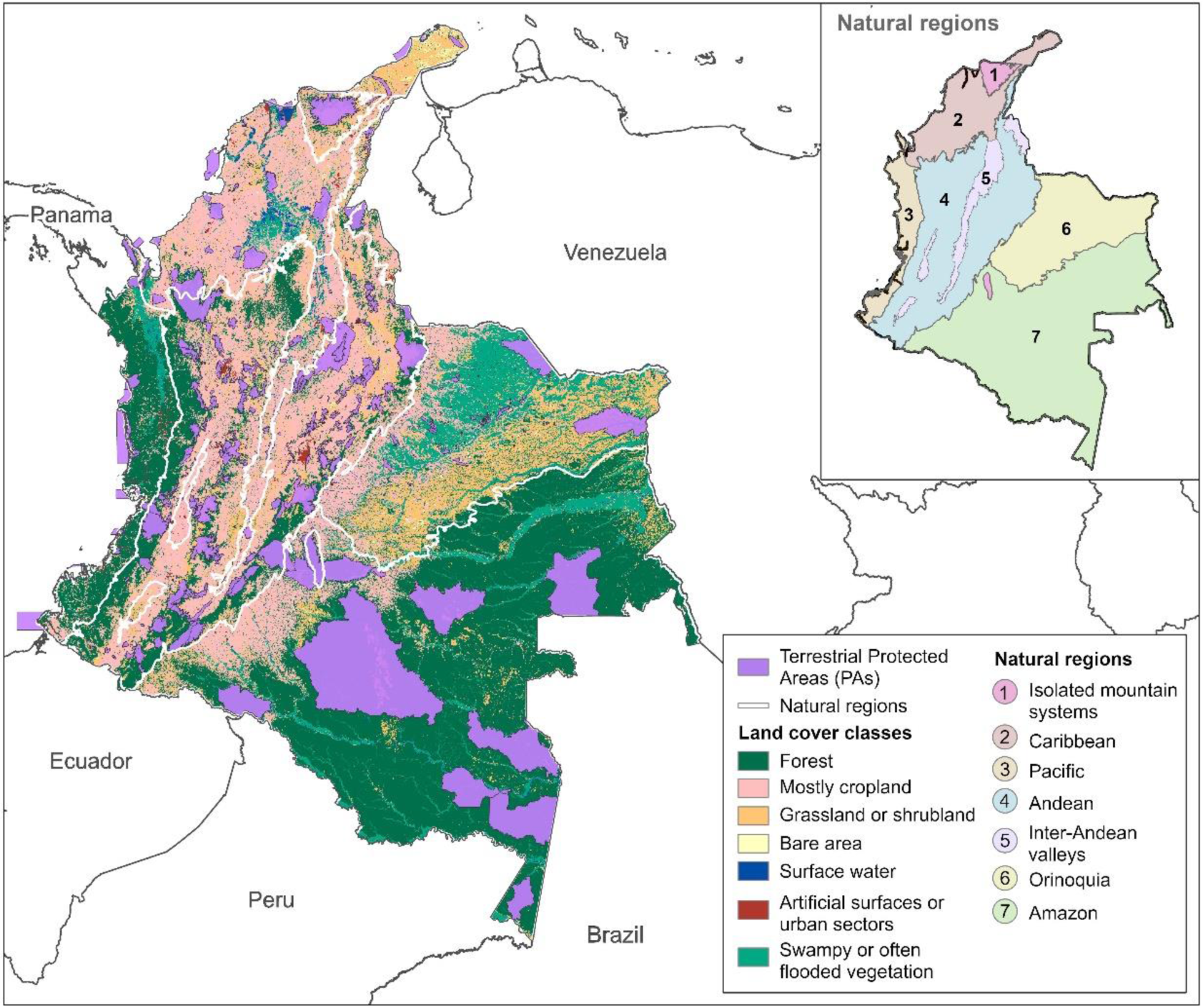
Location of terrestrial PAs belonging to the Colombian National System of Protected Areas (Parques Nacionales Naturales de Colombia, 2022). Different colors indicate the country’s land cover for 2018 (IDEAM, 2021) and regional boundaries.

### Focal bird species

We selected 26 resident and terrestrial bird species (Table 1) representative of natural regions of Colombia (Fig. 1), with a medium to high dependence on natural forest ecosystems according to the IUCN Red List (https://www.iucnredlist.org/). With this selection, we aimed to identify key connectivity areas for multiple species across Colombian forests (Ersoy et al., 2019; Liu et al., 2018). The selection was based on data availability on occurrences, home range sizes for the selected birds or related species of the same genus, and the distribution of each species in Colombia. As information on dispersal distances is sparse, median natal dispersal distances were derived from species home ranges using the relationship proposed by Bowman (2003), which is strong for non-migratory birds. Selected birds were arranged into four groups: small home ranges (<0.2 km^2^) and short dispersal distances (1-5 km); medium home ranges (0.4-10 km^2^) and medium dispersal distances (8-40 km); large home ranges (> 15 km^2^) and long dispersal distances (> 50 km); and a final group including all species.

**Table 1.** Focal bird species selected for the connectivity analyses. Median natal dispersal distances were estimated using the formula proposed by Bowman (2003) and home range values were compiled from the literature (source).

| Common name | Scientific name | Median natal dispersal (km) | Home range (km<sup>2</sup>) | Source |
| --- | --- | --- | --- | --- |
| Berlepsch's Tinamou | *Crypturellus berlepschi* | 4.35 | 0.13 | Estevo et al., 2017 |
| Little Tinamou | *Crypturellus soui* | 3.00 | 0.06 | Guillen et al., 2016 |
| Red-legged Tinamou | *Crypturellus erythropus* | 4.35 | 0.13 | Estevo et al., 2017 |
| Crestless Curassow | *Mitu tomentosum* | 8.75 | 0.53 | Parra et al., 2001; Yumoto, 1999 |
| Chestnut Wood-quail | *Odontophorus hyperythrus* | 2.74 | 0.05 | Franco et al., 2006 |
| Dusky Pigeon | *Patagioenas goodsoni* | 126.55 | 111.21 | Collins et al., 2019; Overton et al., 2010 |
| Great-billed Hermit | *Phaethornis malaris* | 2.41 | 0.04 | Volpe et al., 2016 |
| Ornate Hawk-eagle | *Spizaetus ornatus* | 48.60 | 16.40 | Whitacre, 2012 |
| Black-and-chestnut Eagle | *Spizaetus isidori* | 8.32 | 0.48 | Zuluaga et al., 2018 |
| White Hawk | *Pseudastur albicollis* | 19.71 | 2.69 | Whitacre, 2012 |
| White-throated Hawk | *Buteo albigula* | 59.24 | 24.37 | Llerandi-Roman, 2006 |
| Choco Toucan | *Ramphastos brevis* | 8.03 | 0.45 | Kays et al., 2011; Terborgh et al., 1990 |
| Scaled Piculet | *Picumnus squamulatus* | 2.52 | 0.04 | da Silva et al., 2012 |
| Chestnut Piculet | *Picumnus cinnamomeus* | 2.52 | 0.04 | da Silva et al., 2012 |
| Powerful Woodpecker | *Campephilus pollens* | 9.54 | 0.63 | Ojeda & Chazarreta, 2014 |
| Collared Forest-falcon | *Micrastur semitorquatus* | 36.88 | 9.45 | Thorstrom, 2007 |
| Maroon-tailed Parakeet | *Pyrrhura melanura* | 18.15 | 2.29 | Bianchi, 2010 |
| Scarlet Macaw | *Ara macao* | 144.00 | 144.00 | Brightsmith et al., 2021; Raigoza Figueras, 2014 |
| Yellow-eared Parrot | *Ognorhynchus icterotis* | 100.40 | 70.00 | Salaman et al., 2006 |
| Santa Marta Antpitta | *Grallaria bangsi* | 2.48 | 0.04 | Kattan & Beltran, 2002 |
| Chestnut-naped Antpitta | *Grallaria nuchalis* | 3.66 | 0.09 | Kattan & Beltran, 2002 |
| Yellow-breasted Antpitta | *Grallaria flavotincta* | 2.48 | 0.04 | Kattan & Beltran, 2002 |
| White-bellied Antpitta | *Grallaria hypoleuca* | 2.48 | 0.04 | Kattan & Beltran, 2002 |
| Grey-breasted Wood-wren | *Henicorhina leucophrys* | 2.34 | 0.04 | Vargas et al., 2011 |
| Yellow-crowned Whitestart | *Myioborus flavivertex* | 2.35 | 0.04 | Pomara et al., 2007; Poulsen, 1996 |
| Blue-whiskered Tanager | *Tangara johannae* | 4.93 | 0.17 | Sekercioglu et al., 2007 |

### Species Distribution Models

We developed Species Distribution Models (SDMs) for the focal species to identify their potential suitable habitat across Colombia (Figure 2). First, we obtained geolocated occurrences throughout the complete distribution of each species from the eBird platform (https://ebird.org/home). We only included data recorded after 2000 and up to August 2021 under the sampling protocols specified in eBird as traveling, stationary, and area (The Cornell Lab of Ornithology, 2021). We removed occurrences taken during sampling routes of more than 3 km or 4 hours. We also used the R package CoordinateCleaner (Zizka et al., 2019) to exclude occurrence records with common spatial errors. Additionally, we removed records 10 km beyond published range maps in IUCN and BioModelos (http://biomodelos.humboldt.org.co/). To reduce the effect of sampling bias, occurrences were spatially thinned using the R package spThin (Aiello-Lammens et al., 2015), so there were no records within 5 km of each other. As model predictors, we used the 19 bioclimatic variables from the WorldClim database (Hijmans et al., 2005), with a spatial resolution of approximately 1 km^2^.

**Figure 2.**
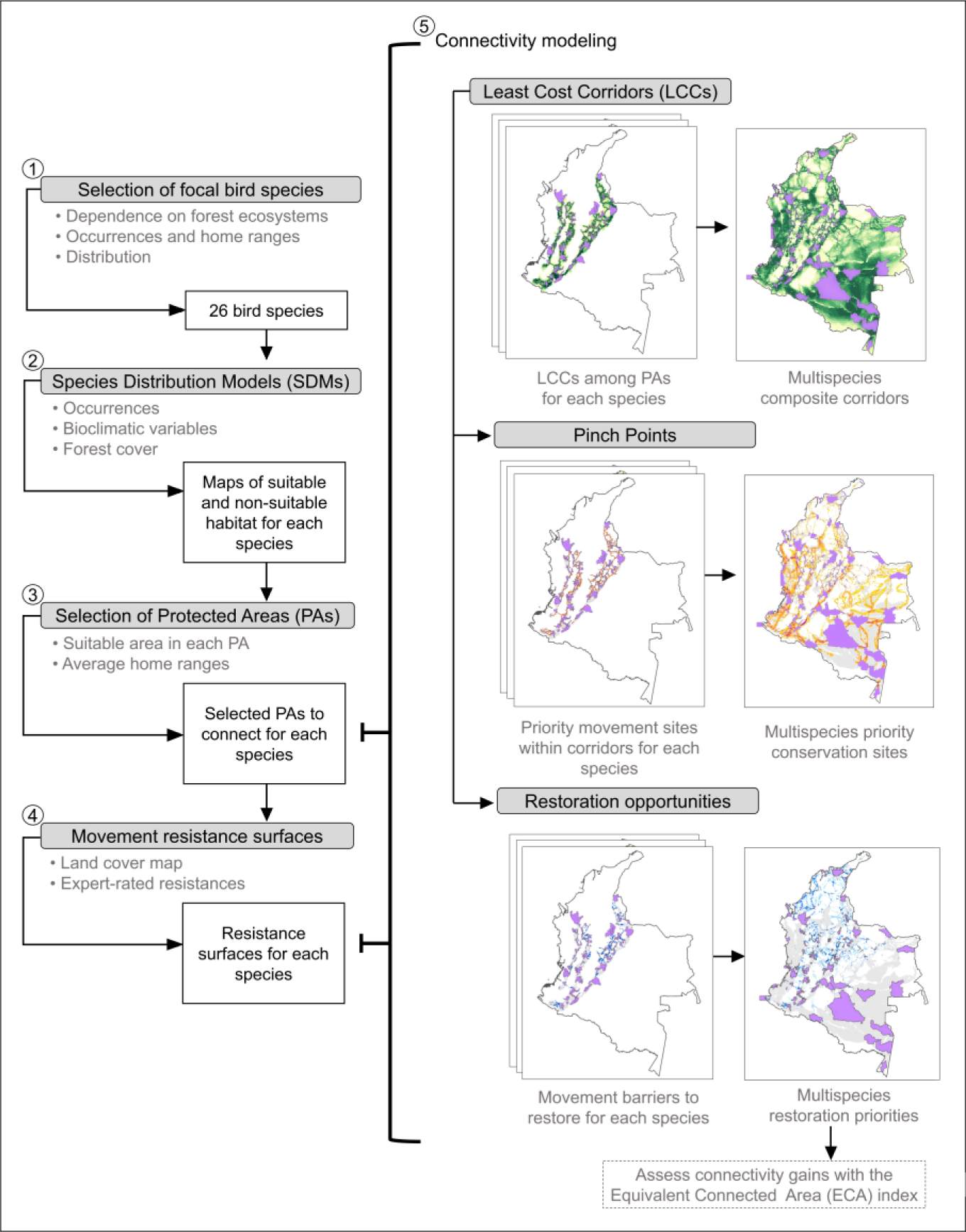
Workflow outlining the methods to identify multispecies corridors among PAs, priority conservation sites, and restoration opportunities.

We modeled the potential habitat for each species using the MaxEnt software (v.3.4.4.; Phillips et al., 2006; Phillips & Dudík, 2008) implemented in the Wallace package (v.1.1.3; Kass et al., 2018). We did not test for collinearity among predictors because Maxent models can regulate redundant variables through a regularization parameter optimized in ENMeval (Elith et al., 2011; Merow et al., 2013). The accessible area or M of the models was defined as the minimum convex polygon of the thinned occurrences for each species plus a 10 km buffer. Models were run with 10,000 randomly sampled background points, and based on the number of occurrences, we partitioned the data using the jackknife (<25 records), checkerboard1 (>30 records), or checkerboard2 (>100 records) methods (Muscarella et al., 2014). Additionally, we tested a Target-Group Background selection (TGB) approach to account for sampling bias in the models (Phillips et al., 2009). To implement TGB, we used the complete set of cleaned records of all focal birds as background points for each species. We selected regularization multipliers from 0.5 to 4 with 0.5 intervals to test the following feature class combinations: Linear (L), Linear Quadratic (LQ), Hinge (H), and Linear Quadratic Hinge (LQH). We assessed the optimal parameters of the resulting models based on the lowest average test omission rate (10^th^ percentile) followed by the highest average test AUC (Shcheglovitova & Anderson, 2013). We used the selected settings to generate habitat suitability predictions across Colombia.

Final cloglog predictions for each species were converted to binary maps of suitable/non-suitable habitat using the 10^th^ percentile training presence threshold. Model selection between random background points and TGB approaches was performed by JV through visual comparison with field guides and on-the-ground knowledge of bird distributions (Bateman et al., 2020; Velásquez-Tibatá et al., 2019). Finally, we used the 2018 national land cover map (IDEAM, 2021) to reclassify cells with less than 50% natural forest cover in 1 km^2^ as unsuitable.

### Protected areas and resistance surfaces

With the distributions of potential suitable habitat and home range estimates, we selected the PAs to include in the connectivity analysis of each species (Figure 2). To avoid mapping errors near range limits, we buffered the species distributions to achieve a ∼20% increase in their area (Koen et al., 2014; Wade et al., 2015). Then, we selected the PAs having a suitable area greater than or equal to the average home range size reported in the literature for each species (Table 1). We considered all PAs within Colombia’s National System of Protected Areas (SINAP), including national, subnational, public, and private PAs (Figure 1).

Next, we built the resistance surfaces to represent the degree to which different land covers impede or facilitate the movement of each species across its distribution (Figure 2; Taylor et al., 1993; Zeller et al., 2012). Since empirical data on the movement ecology of birds in Colombia are limited, we consulted 17 ornithology experts to estimate the resistance of different land covers. Each expert completed an electronic form (Supplementary Material Fig. A1) estimating the difficulty of crossing at least 300 m through 7 different land categories for each species they were familiar with. Land cover classes were obtained from the Colombian land cover map for 2018 (300 m resolution; IDEAM, 2021) and generalized into the following categories: forest, mostly cropland, grassland or shrubland, artificial surfaces or urban sectors, bare areas, swampy or often flooded vegetation, and surface water (Figure 1). Experts ranked each class using a Likert scale (Supplementary Material Fig. A1) ranging from absolute resistance (100) to no resistance to movement (0). The average number of experts who filled out the form per species was 5 (SD = 3), and only two species, *Phaethornis malaris* and *Pseudastur albicollis*, had a single entry. Finally, we used the experts’ responses to create the resistance surfaces using the average values for each land cover class and species (Ersoy et al., 2019; Supplementary Material Fig. A2).

### Connectivity modeling

We used each species’ PAs and resistance surfaces as inputs in Linkage Mapper (McRae & Kavanagh, 2011), an ArcGIS toolbox for performing connectivity analyses. Firstly, we used the Linkage Pathways tool to map the Least-Cost Corridors (LCCs) among PAs (Figure 2). Linkage Pathways produces rasters of cost-weighted distance (CWD) values between pairs of neighboring PAs, highlighting the path with the lowest cumulative movement resistance or least-cost path (LCP; Adriaensen et al., 2003; McRae & Kavanagh, 2021). It then normalizes and mosaics the results from each pair of PAs, creating a single composite showing all LCPs and LCCs, which are wider swathes of pixels having slightly higher movement resistance than the optimal path (McRae & Kavanagh, 2021; Schrott & Shinn, 2020). We ran the tool dropping LCPs intersecting intermediate PAs and allowing a maximum of four connected nearest neighbors. We modeled LCCs regardless of length due to the uncertainty surrounding our dispersal distance estimates and because they may represent opportunities for management actions to enhance connectivity (Schrott & Shinn, 2020).

We created four composite maps combining the results for all species and the three groups based on home range size (Figure 2). To do this, we reclassified the corridor rasters for each species so that the lowest-cost decile had a value of 10, the next decile a value of 9, and so on, assigning higher values to lower-cost pixels (Belote et al., 2016). Then, we added the reclassified rasters from each group and divided them by the number of species distributed in each cell. In this way, the group maps range from 1 to 10, with 10 representing the corridors having the lowest cost-distance values for all the species distributed there.

We used the Pinchpoint Mapper tool of Linkage Mapper to identify priority conservation sites within the identified corridors (Figure 2). Pinchpoint Mapper applies circuit theory by running Circuitscape (McRae et al., 2008) within the LCCs to produce current flow maps showing the net probabilities of passing through each cell as birds move between PAs. Therefore, pinch points are places with high current flow concentrations indicating a greater movement probability or the lack of alternative paths to move between PAs (McRae et al., 2008). Loss of suitable habitat at pinch points could affect connectivity significantly (Castilho et al., 2015; McRae et al., 2008). To run the tool, we tested 1, 5, and 10 km as corridors’ width limits in CWD units. We selected the 10 km width to reflect the uncertainty in the connectivity models and used the adjacent pair method. To produce the multispecies results, we reclassified the individual species maps to zeros and ones if the current values were below or above the 50th percentile, respectively (González-Saucedo et al., 2021). We summed the individual maps for each of the four species groups so that the pixel values indicate the number of species with overlapping pinch points.

Areas that could improve connectivity among PAs through restoration were identified using Barrier Mapper (Figure 2). This tool identifies movement barriers significantly impacting the quality and location of corridors that when restored, provide the highest increase in connectivity (McRae et al., 2012). Barriers are features in the landscape matrix that hinder movement between habitat patches and could be, for example, land covers with high movement resistance around PAs. Barrier Mapper applies a moving window across the resistance surfaces, simulating pixel restoration within the window. It estimates the restoration potential or improvement score as the corridors’ least-cost distance reduction (McRae et al., 2012). We applied moving windows of 300, 600, and 900 m based on the size of Colombia’s small, medium, and large-scale restoration projects (Murcia & Guariguata, 2014). We combined the results of all radios into a single map for each species using the maximum improvement scores. To produce the multispecies composites, we reclassified the species scores to zero if they were below the 50th percentile and one if they were higher (González-Saucedo et al., 2021). We then added the rasters for the four species groups to obtain maps with pixel values indicating how many species had a barrier that, if restored, would have an improvement score above the 50th percentile. Subsequently, we excluded natural barriers, such as water bodies, from anthropic barriers with restoration potential under the classes of mostly cropland, grassland or shrubland, artificial surface or urban sectors, and bare area. Using the 2014 conversion map of potential natural ecosystems developed by Etter et al. (2020b), we excluded natural grasslands, shrublands, and sandy lands from the restoration results.

We estimated the potential increase in connectivity among PAs by restoring priority sites using the Equivalent Connected Area (ECA) index, which integrates intra-patch and inter-patch connectivity (Saura et al., 2011; Saura & Pascual-Hortal, 2007). The ECA index is defined as the area of a single continuous patch that would provide the same probability of connectivity value as the actual landscape pattern within an area of interest (Saura et al., 2011). Using the Makurhini R package (Godínez-Gómez & Correa Ayram, 2020) we calculated the percentage of variation of ECA (dECA) and habitat area (dA) under two scenarios. The current scenario included the natural forest cover reported for Colombia in 2018 (IDEAM, 2021), while the second added the forest cover derived from restoration. For these calculations, we used the Powerful Woodpecker (*Campephilus pollens*), as a representative species from our study group as it has a mean home range of 0.6 km^2^ and a median dispersal distance of 9.5 km (dispersal probability = 0.5). This distance of approximately 10 km is commonly used in studies to estimate changes in landscape connectivity because it is a central estimate for the range of dispersal distances among terrestrial vertebrates (Castillo et al., 2020; Murillo-Sandoval et al., 2022; Saura et al., 2019).

## Results

### Corridors among PAs

Based on responses from bird experts, we found that forests had the lowest resistance value for all species, while artificial surfaces presented the highest resistance to movement in most cases (Supplementary Material Fig. A3). Based on our Linkage Pathways analysis, the composite map of the small home range group included the highest number of PAs (666) and LCCs, while the large home range group had the least amount of LCCs and included the fewest PAs (191; Figure 3B).

**Figure 3.**
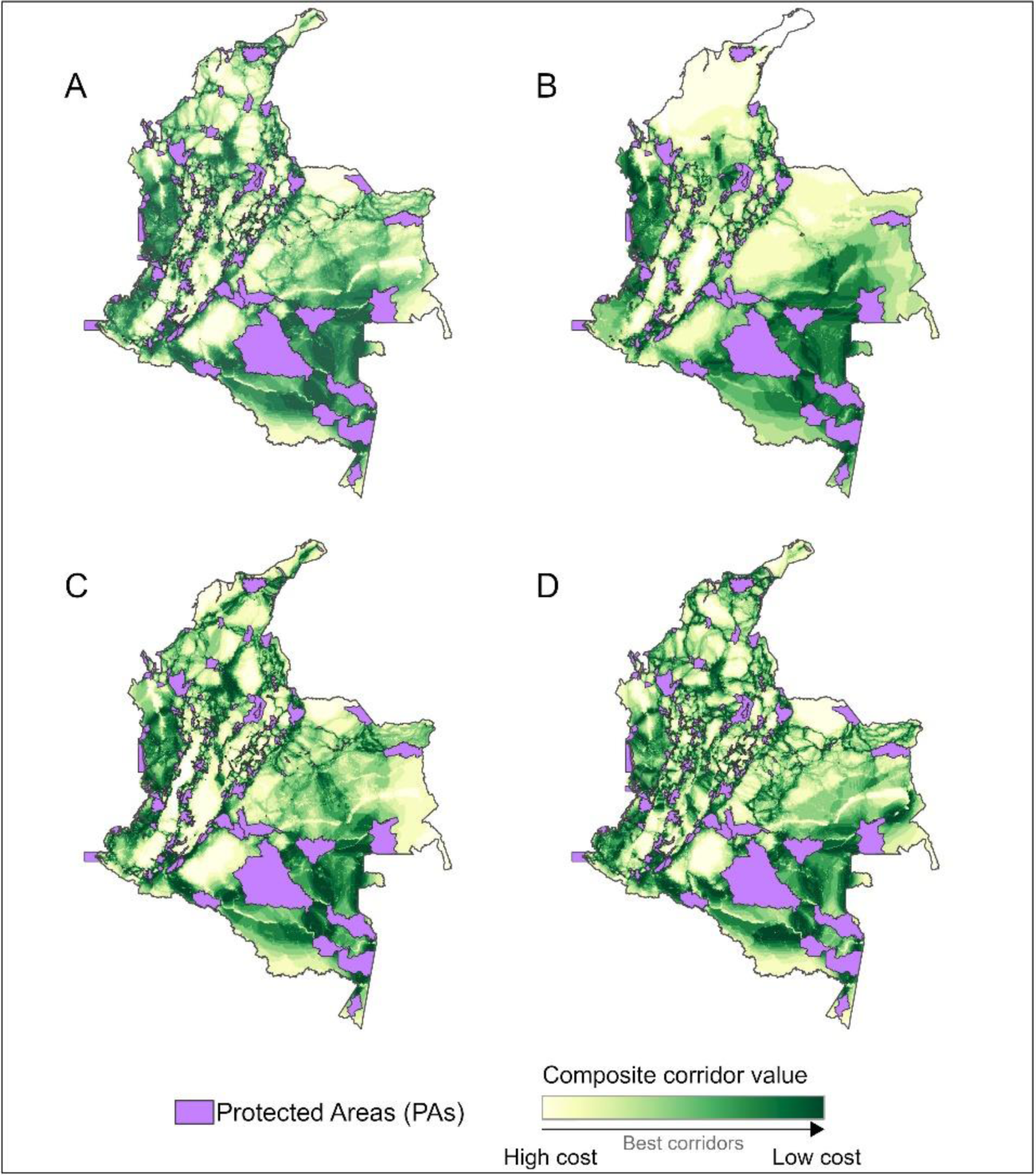
Composite maps showing the least-cost corridors for four different species groups. All 26 focal species (A). Birds with large home ranges (> 15 km^2^; B). Birds with medium home ranges (0.4-10 km^2^; C). Birds with small home ranges (<0.2 km^2^; D). Greenest areas indicate pixels having the lowest cost-distance values for all the species distributed there.

Hereafter we will focus on the composite map for all species summarizing the results for all home range groups (Figure 3A). This composite includes 723 PAs, and the corridors cover 229,205 km^2^ under the top 20% composite values and 581,531 km^2^ under the top 50%. Forests dominate the extent of the corridors, covering 85% and 68% of the area under the top 20% and 50% values, respectively. However, these results vary by geographic region (Figure 1): croplands exceed forests in the LCCs of the Caribbean and Inter-Andean Valleys regions (Supplementary Table B1). Conversely, the Amazon, Andes, and Pacific regions have the greatest forest cover within their LCCs (Supplementary Table B1).

### Conservation priorities

Based on the pinch-point analysis, we prioritized 212,769 km^2^ that are critical to maintaining connectivity among PAs for our study species (Figure 4). These priorities consist mainly of forested lands (69%) and are distributed predominantly across the Andean (36%) and Amazon (31%) regions (Supplementary Table B2). Some areas stand out as important to maintain multi-species movement, such as the corridor between the Páramo El Atravesado Reserve (IUCN category VI) and the Bosque de Los Guayupes Regional Park (IUCN category II; Figure 4A). In the Amazon, pinch points constitute more restricted strips within the wide LCCs of the region, as observed in the Andean-Amazonian transition with the corridor between La Paya and Serranía de los Churumbelos Auka-Wasi National Parks (IUCN category II; Figure 4B).

**Figure 4.**
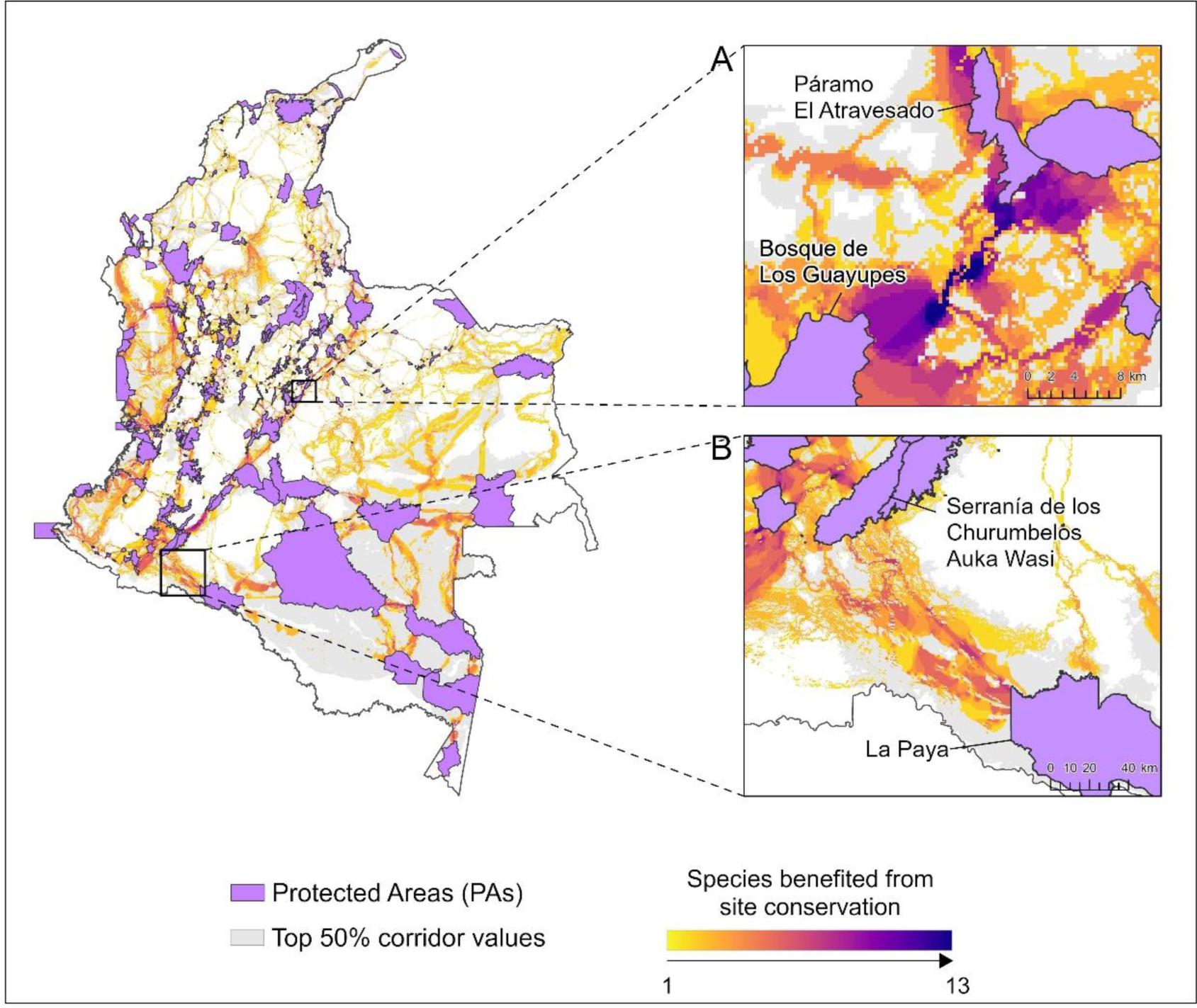
Priority conservation sites (pinch points) within the Least Cost Corridors (LCCs) for all study species. Inset A, important sites for the movement conservation of multiple species across the corridor between the Páramo El Atravesado Reserve and Bosque de Los Guayupes Regional Park. Inset B, key connectivity sites within the corridor between the La Paya and Serranía de los Churumbelos Auka Wasi National Parks. Dark purple hues indicate pixels where more species had net movement probabilities above the 50th percentile.

### Restoration priorities

We identified 79,228 km^2^ with the potential to improve PAs connectivity by being restored to forests (Figure 5). These areas comprise mainly croplands (78%) and non-native grasslands and shrublands (20%). Restoration priorities concentrate in the Andes (45%) and the Caribbean (29%) regions (Supplementary Table B3). While most restoration priorities concentrate on narrow corridors connecting pairs of PAs in some areas they extend to the matrix and connect multiple PAs. For example, the network of Civil Society Reserves (IUCN category VI) surrounding the Los Colorados Fauna & Flora Sanctuary (IUCN category IV) in the Caribbean region is embedded in a matrix of crops that, if restored, could benefit up to 5 species (Figure 5A). A similar situation occurs in a cluster of reserves across an altitudinal gradient in the Western Andes (Figure 5B; IUCN category VI). Our ECA analysis indicates that restoring the priorities identified could increase the national forest cover by 7% and functional connectivity by 14% (Figure 6). Thus, the increase in connected habitat measured by the ECA index is more substantial than the increase in total habitat area (dECA>dA>0).

**Figure 5.**
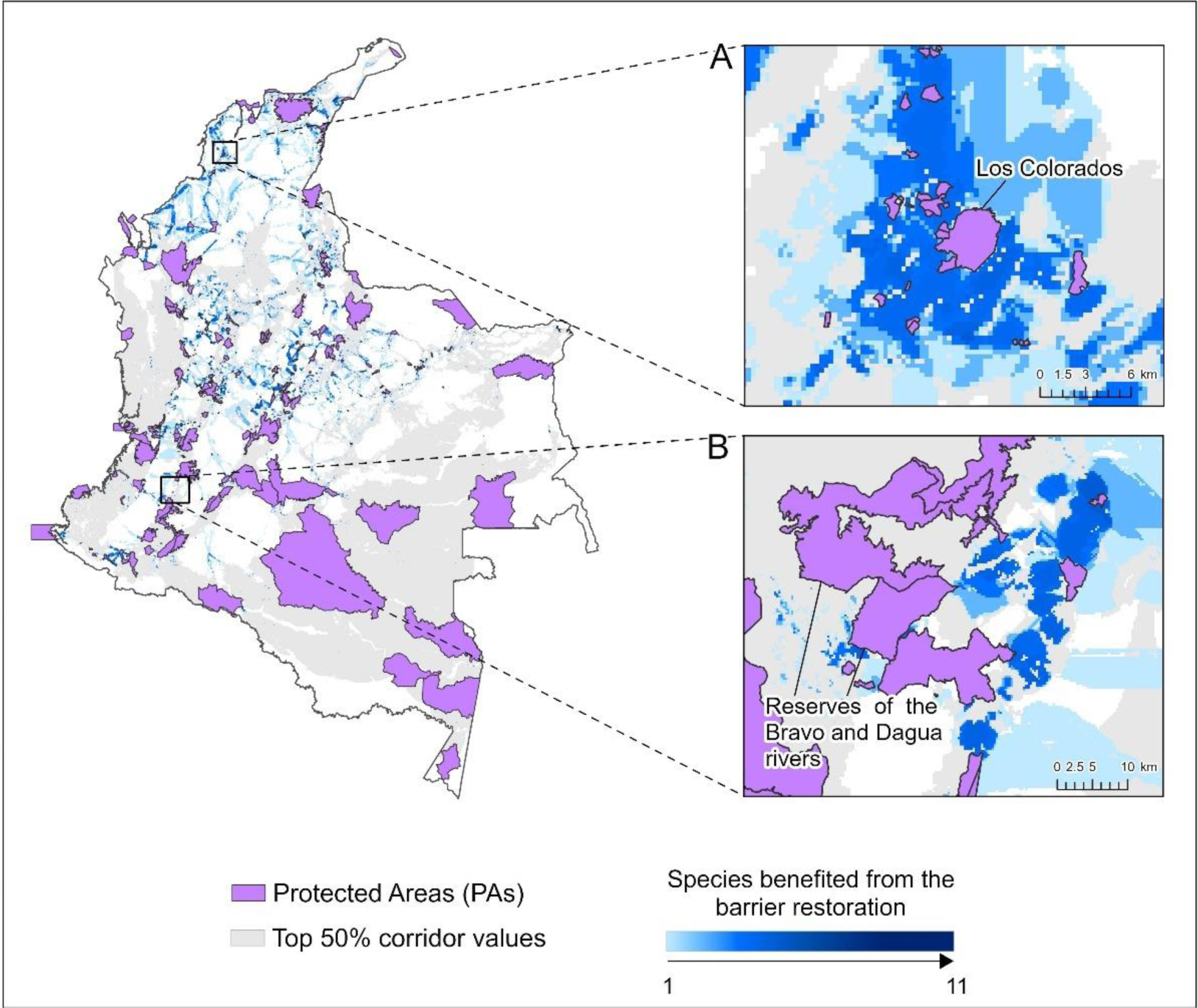
Restoration priorities to improve PA connectivity. A) Restoration priorities around the Los Colorados Flora and Fauna Sanctuary (A). Restoration priorities surrounding the Regional Forest Reserves of the Bravo and Dagua rivers (B). Increasingly dark shades of blue indicate pixels where more species benefit from restoration.

**Figure 6.**
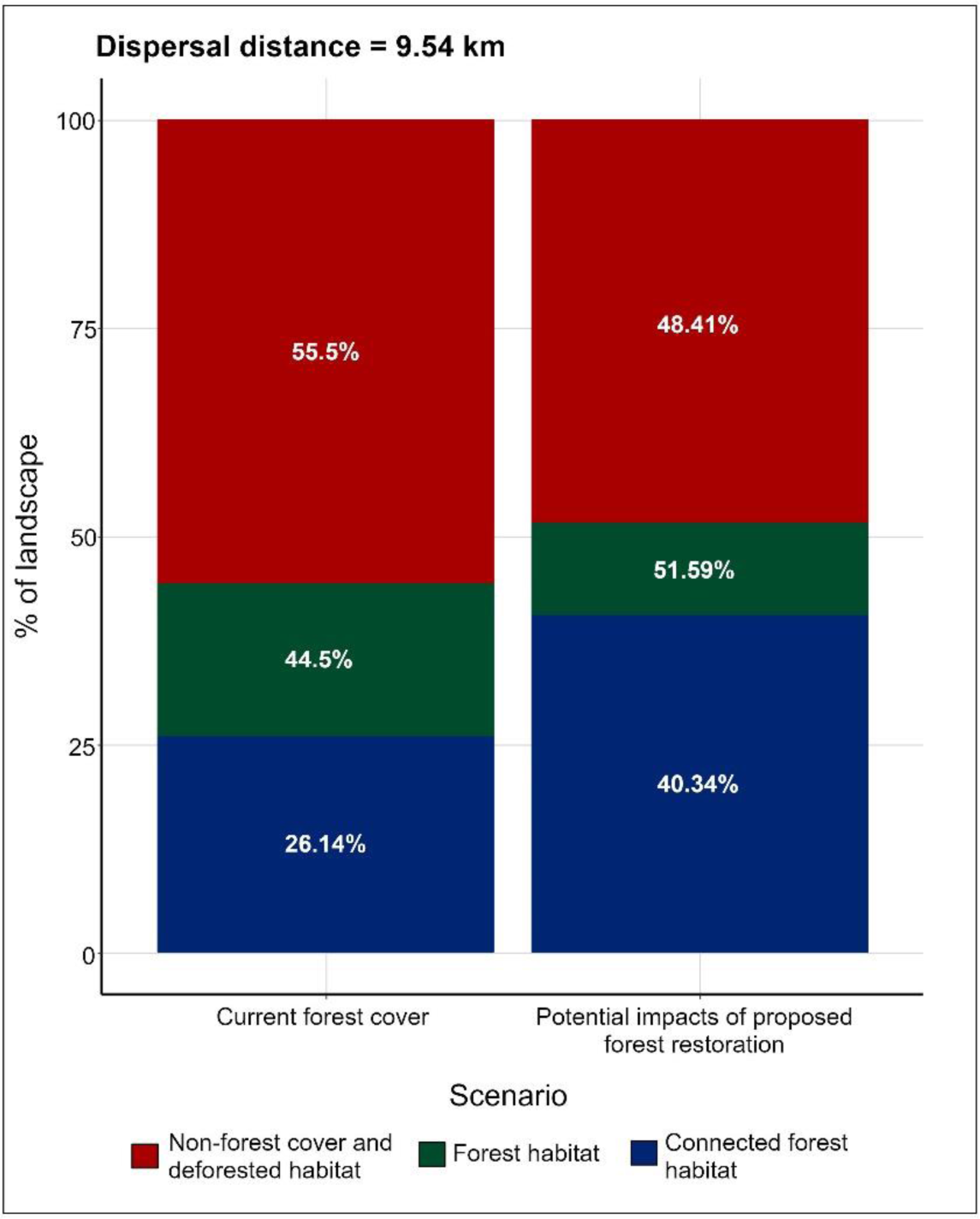
Estimated gain in functional connectivity and forest cover derived from the restoration of the identified priorities according to the Equivalent Connected Area (ECA) index changes.

## Discussion

The expansion and intensification of human activities have led to the rapid loss and fragmentation of tropical forests (Barlow et al., 2016; Correa Ayram et al., 2020; Hoang & Kanemoto, 2021). PAs currently constitute a central strategy to preserve remnants of continuous natural forests, and preserving ecological connections between them is vital for species and ecosystem conservation (Hilty et al., 2020; Watson et al., 2014). Here, we identified corridors primarily composed of natural forests, representing potential ecological linkages among national and subnational PAs. Through a combination of least-cost and circuit-theory models, our study prioritized the protection of 212,551 km^2^ and restoration of 79,203 km^2^ to maintain and enhance connectivity among PAs. This study is the first to assess bird functional connectivity for the Colombian PA network and it could aid in guiding the country’s effort in safeguarding its exceptional biodiversity under global change.

Our results agree partially with corridors identified for other taxonomic groups in Colombia. For instance, the corridors we identified in the Caribbean region between the Sierra Nevada de Santa Marta National Park and the Serranía del Perijá, as well as between the Catatumbo Bari and Tama National Parks, are essential for the movements of jaguars and other carnivorous animals (Payán-Garrido & von Hildebrand, 2016; Zárrate Charry et al., 2022). Also, our results are strikingly similar to Areiza et al.’s (2019) and Pineda’s (2022) for the Andean and Amazon regions, revealing comparable connections between the Farallones de Cali and Tatamá National Parks in the western Andes and between the Chiribiquete National Park and other Amazonian PAs such as La Paya, Tinigua, and Nukak. These similarities reinforce the ecological importance of these corridors for a wide range of species and strengthen the case for their inclusion in conservation planning processes in Colombia. Additionally, this study substantially complements existing connectivity prioritization efforts, extending them to consider bird functional connectivity across the different natural regions of the country.

Regarding restoration opportunities, our findings partially align with existing national priorities, such as the National Restoration Plan (Vanegas Pinzón et al., 2015) and the restoration priorities for high-risk ecosystems in Colombia (Etter et al., 2020a). Significant overlap exists between these plans and our results in the eastern Andes deforested areas (from the departments of Cundinamarca to Norte de Santander) and the Caribbean region (Antioquia to Cesar). However, 82% of our restoration priorities are currently unaddressed by current restoration plans, highlighting that restoration may not always result in improved ecological connectivity. Moreover, the connectivity gain resulting from the restoration priorities we identified was double the amount of gain in forest area. Therefore, our results suggest that targeted restoration of small crucial sites could be more effective than restoring large areas with limited potential for enhancing connectivity.

Ecological connectivity research should be ideally based on actual species movement data derived from GPS tags, telemetry studies, or landscape genetics (Epps et al., 2007; Sawyer et al., 2011; Zeller et al., 2018). These data are also the most reliable approach for validating connectivity models and their predictions of movement routes (Pullinger & Johnson, 2010; Sawyer et al., 2011). We used expert criteria to develop resistance surfaces due to the limited availability of movement data in the tropics (Pillay et al., 2022; Robinson et al., 2018). While this has frequently been criticized as subjective (Zeller et al., 2012), research indicates that resistance surfaces based on expert-assigned scores can yield results comparable, at a coarse scale, to those derived from empirical data (Keeley et al., 2016; Liu et al., 2018; Pither et al., 2023). Additionally, while a substantial portion of independent occurrences of the focal species aligned with the high connectivity values and selected PAs from our study (Supplementary Table C1), field research is still needed to validate resistance surfaces and assess the effectiveness of corridors in facilitating movement for our study species (Gregory & Beier, 2014; Montealegre-Talero et al., 2017). Despite these shortcomings, we believe our study presents a sensible approach to identifying functional forest corridors for a representative sample of Colombian birds under current data limitations and, as such, is a valuable step towards including functional connectivity as part of Colombia’s conservation agenda that could be improved as movement studies proliferate by harnessing new technologies such as the MOTUS network.

### Conservation implications

This study aims to provide decision-makers and planning institutions in Colombia with spatial guidance for conserving and restoring connectivity among PAs, which can benefit birds and other forest-dependent species. This information is highly relevant considering that natural areas occupy less than half of the country’s land, and human impacts on ecosystems have increased substantially in the last decades (Correa Ayram et al., 2020; González et al., 2018). The most recent National Deforestation Policy warns that approximately 2.8 million hectares of forest have been cleared between 2000 and 2019, primarily due to the expansion of extensive cattle ranching and small to medium-scale agriculture (Consejo Nacional de Política Económica y Social, 2020; González et al., 2018). Projections indicate that by 2030, areas with a high diversity of threatened, endemic, and forest-specialist birds may experience increased human pressure and habitat transformation (Ocampo-Peñuela et al., 2022). Therefore, it is vital to allocate resources towards the conservation and restoration of priority sites, as identified in this study, for achieving effective 30×30 and post-2020 national and global targets (Convention on Biological Diversity, 2021; Gobierno de Colombia, 2021; Oppler et al., 2021).

Our analysis helps to identify and delineate corridors and sites allowing bird movement among PAs that may be crucial for multiple biological processes. Hence, our results could help conservation practitioners and regional planning organizations implement different strategies, including payments for environmental services (PES), new PAs, and other effective area-based conservation measures (OECMs). Implementing these strategies is urgent due to the growing deforestation risk in critical areas that allow and will allow the movement of species under climate change and increased human pressure. For example, the Andean-Amazonian Transition Belt has experienced a surge in deforestation since the signing of the peace agreement between the National Government and the Revolutionary Armed Forces of Colombia (FARC) in 2016 (Clerici et al., 2020; Murillo-Sandoval et al., 2022, 2023). Due to the resultant increase in connected forest loss (Murillo-Sandoval et al., 2022), safeguarding the different corridors we identified in this area, such as the one between La Paya and Serranía de los Churumbelos Auka Wasi National Parks (Figure 4B), could be crucial for species migration along the Amazon-Andes elevation gradient. Critical linkages should also be conserved across the Serranía del Darién and Northern Andes, around Los Katios and Paramillo National Parks, to maintain the functional connectivity between Meso and South America (Correa Ayram et al., 2019, 2020).

Although protecting critical connectivity linkages is essential for biodiversity conservation, ecological restoration of transformed areas is also required to face the threats of climate change and forest loss and fragmentation (Aronson & Alexander, 2013; Etter et al., 2020a; Masson-Delmotte et al., 2018; Perring et al., 2018). Our analysis shows that with a ∼7% increase in national forest cover, mainly in the Caribbean and Andean Mountain ranges and valleys, it is possible to gain twice the percentage of functional connectivity for birds with dispersal distances greater than 9 km. The urgent need for restoration in the Andes and Caribbean has been pointed out on multiple occasions because these regions are the most transformed in the country despite being home to a great diversity of ecosystems and species, including the critically endangered tropical dry forest biome (Aldana-Domínguez et al., 2017; Etter, Andrade, Saavedra, et al., 2017; Etter et al., 2020a; Etter & van Wyngaarden, 2000). Restoration in these regions could lead to significant conservation benefits for birds, as it helps to mitigate the impacts of climate change and to connect remnant habitat fragments (Masson-Delmotte et al., 2018; Montealegre-Talero et al., 2017; Torres et al., 2022).

Despite the importance of ecological restoration, numerous challenges arise due to its high implementation costs and tradeoff with productive activities in established agricultural landscapes (Brancalion et al., 2019; Tanentzap, 2015; Vergara et al., 2016). However, alternative actions such as planting isolated trees in agricultural lands, live fences, sylviculture, and protection of riparian forests hold the potential to improve species’ ability to move across agroecosystems (Díaz-Bohórquez et al., 2014; García-Núñez et al., 2020; Harvey et al., 2004; Lentijo et al., 2022; Martínez-Salinas & DeClerck, 2021; Simioni et al., 2022). Future research is needed to explore the potential of these management actions to restore and enhance the connectivity of natural habitats at large spatial scales.

## Conclusion

The movement of birds between protected areas is vital to ensure their long-term persistence and guarantee the provision of the ecosystem services they offer. Given the historical and anticipated impacts of forest loss and fragmentation in Colombia, securing the protection and restoration of critical areas for connectivity is imperative. By identifying corridors among national and subnational PAs and critical sites to enhance and preserve species movements, we have provided spatial guidance to effectively guide conservation, restoration, and sustainable management efforts to secure an interconnected system of PAs in Colombia. Notably, this study’s emphasis on bird functional connectivity adds a new dimension to Colombia’s conservation planning efforts currently being used to direct investments for new protected areas through the Conserva Aves initiative (https://conservaves.redlac.org/). In this way, our study contributes significantly to the mission of the National Strategy for the Conservation of Birds to strengthen the conservation of birds in the country and to attain ambitious national and global environmental goals by 2030 and beyond.

## Acknowledgments

We want to express our sincere gratitude to the following people who generously shared their time, knowledge, and expertise with us: Alejandro Pinto, Diego Calderón-Franco, Diego Carantón, Felipe Estela, Gabriel Utria, Giovanni Cárdenas, Gloria Lentijo, Gustavo Londoño, Héctor Rivera, Jorge Muñoz, Manuel Espejo Hoyos, Nick Bayly, Noemi Moreno Salazar, Paola Campo Soto, Pedro Arturo Camargo Martínez, and Sergio Chaparro Herrera. Financial support to JVT was provided by NASA Award 80NSSC21K1943 - Ecological Conservation Program. We are also grateful to the Microsoft AI for Earth grant for providing us with the computational tools to analyze and process our data.

## Data Availability

Results are available in GeoTIFF format through https://doi.org/10.5281/zenodo.7947108

