## Supplementary Material A for "Prioritizing ecological connectivity among protected areas in Colombia using a functional approach for birds"

**Project Description**

The objective of this project is to identify the most critical sites for conserving and restoring connectivity among protected areas in Colombia. For its development, we have selected 26 focal bird species, for which we will delineate potential ecological corridors among protected areas based on a land cover map from the year 2020 (resolution of 300 meters). The purpose of this form is to assign a resistance value to different types of land cover.

Please provide your general information and select the species for which you will assign resistance values.

### Date

### Name

### Institution

### Role

### Selected species

Please estimate the relative difficulty for the species to move at least 300 meters through different land cover types. On this scale, "No resistance" represents land cover types that the species can traverse without restriction. The option "Moderate resistance" indicates that it is a land cover that the species occasionally crosses to reach more suitable habitat patches. Land cover with "Absolute resistance" is a complete physical barrier to movement, meaning the species never crosses it.

### Mostly Agriculture

Includes herbaceous, shrub, and tree crops. Also, mosaics of crops (>50%) with natural vegetation (<50%).

Absolute resistance — High resistance — Moderate resistance — Low resistance — No resistance

**Grasslands and Shrubs**

| Absolute resistance | High resistance | Moderate resistance | Low resistance | No resistance |
|---|---|---|---|---|

**Forests**

Includes 300-meter resolution cells with at least 15% non-flooded forest cover.

| Absolute resistance | High resistance | Moderate resistance | Low resistance | No resistance |
|---|---|---|---|---|

**Sparse Vegetation**

Sparse vegetation (trees, shrubs, and herbaceous cover < 15%).

| Absolute resistance | High resistance | Moderate resistance | Low resistance | No resistance |
|---|---|---|---|---|

**Urban**

Artificial surfaces or urban areas.

| Absolute resistance | High resistance | Moderate resistance | Low resistance | No resistance |
|---|---|---|---|---|

**Bare Areas**

Rock outcrops or bare soil.

| Absolute resistance | High resistance | Moderate resistance | Low resistance | No resistance |
|---|---|---|---|---|

**Swampy or Frequently Flooded Vegetation**

Tree, shrub, or herbaceous cover flooded by fresh/saline/brackish water.

| Absolute resistance | High resistance | Moderate resistance | Low resistance | No resistance |
|---|---|---|---|---|

**Water Bodies**

Rivers, lagoons, and reservoirs at least 300 meters wide.

| Absolute resistance | High resistance | Moderate resistance | Low resistance | No resistance |
|---|---|---|---|---|

Figure A1. Example of the structure of the electronic form used to gather expert ornithological estimates on the difficulty of focal bird species in crossing various land cover types.

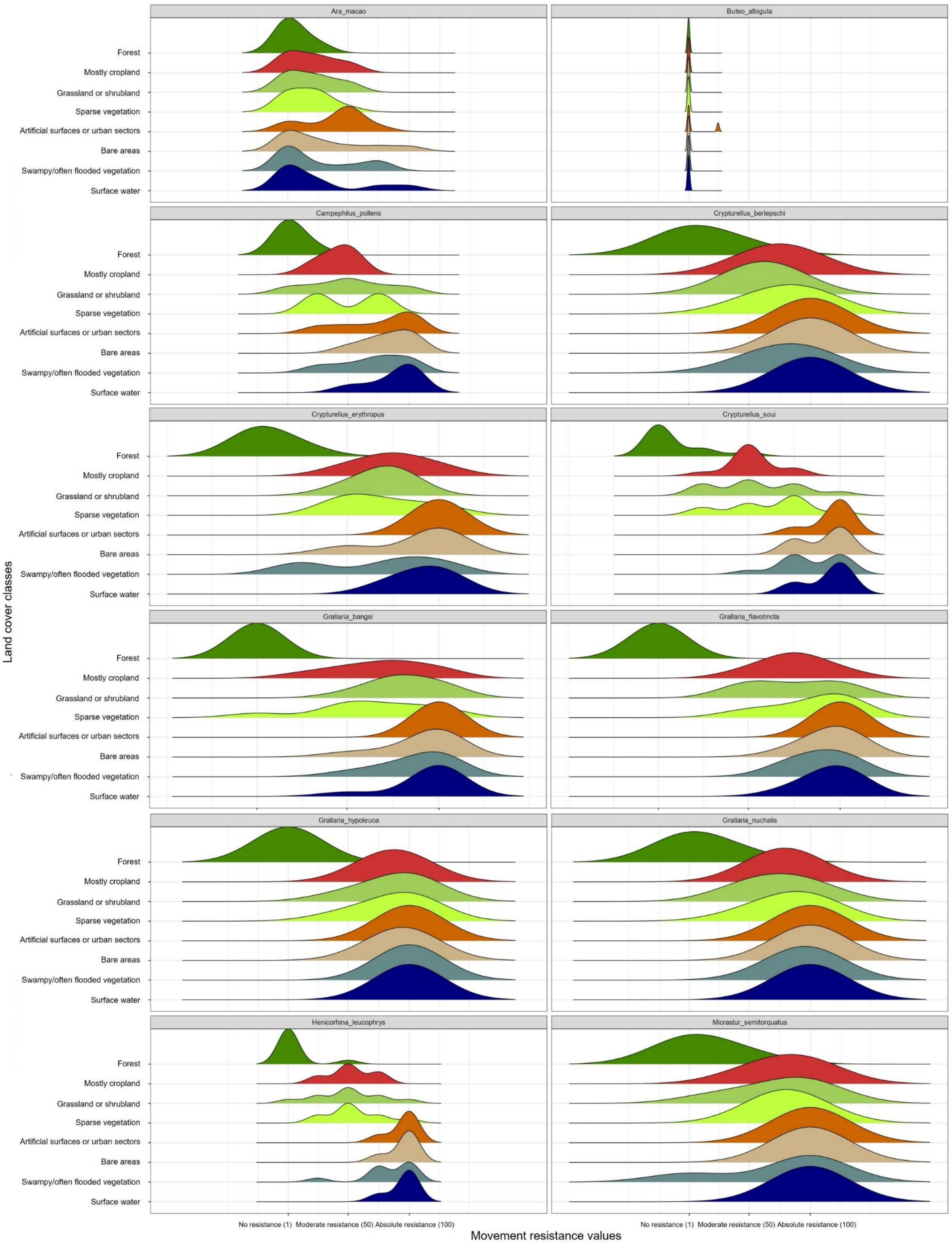
Ara_macao
Buteo_albigula
Campephilus_pollens
Crypturellus_berlepschi
Crypturellus_erythropus
Crypturellus_soui
Grallaria_bangsi
Grallaria_flavotincta
Grallaria_hypoleuca
Grallaria_nuchalis
Henicorhina_leucophrys
Micrastur_semitorquatus
Forest
Mostly cropland
Grassland or shrubland
Sparse vegetation
Artificial surfaces or urban sectors
Bare areas
Swampy/often flooded vegetation
Surface water
Land cover classes
No resistance (1) Moderate resistance (50) Absolute resistance (100)
Movement resistance values

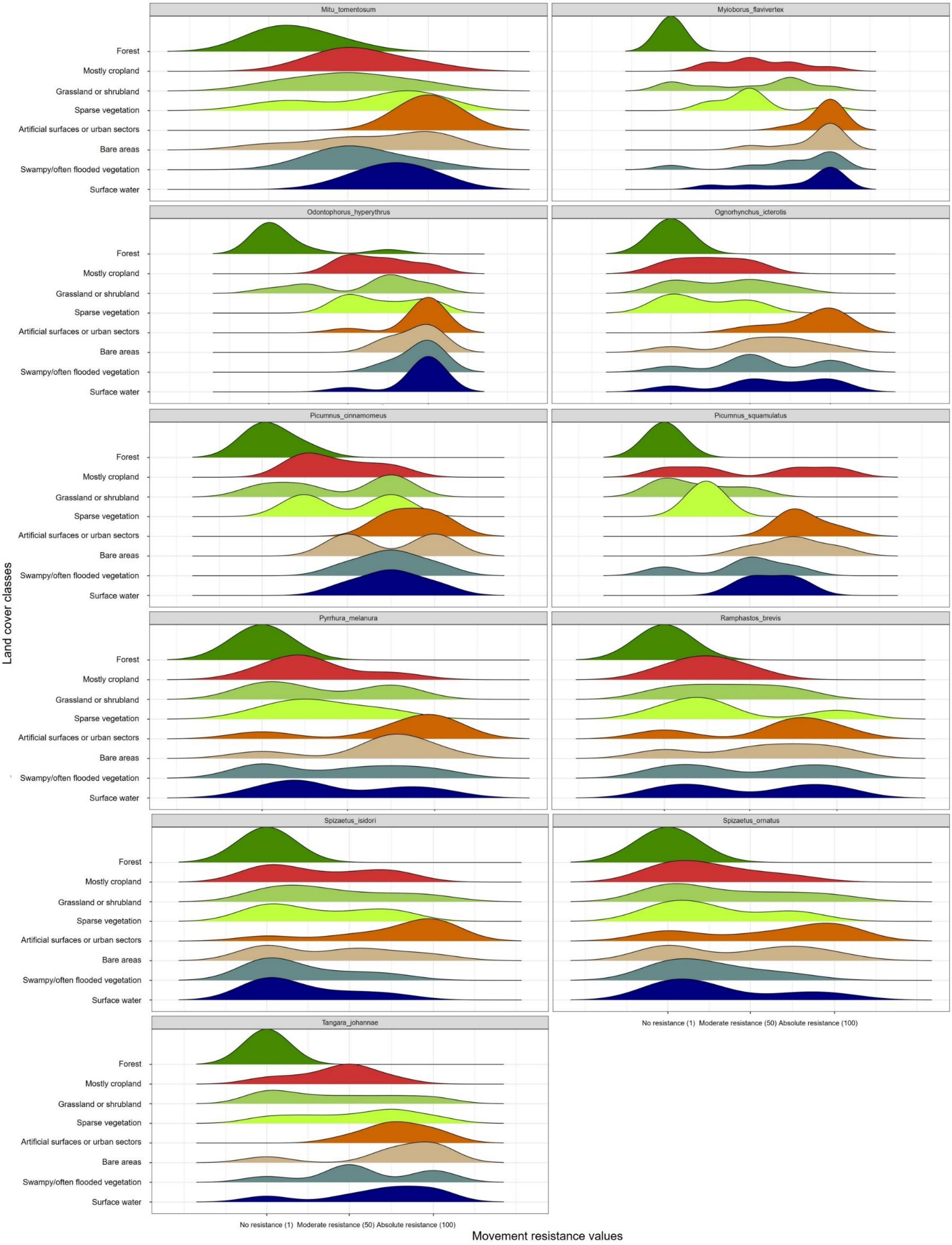


Figure A2. Distribution of expert-rated resistance to movement values for each of the 26 focal bird species.

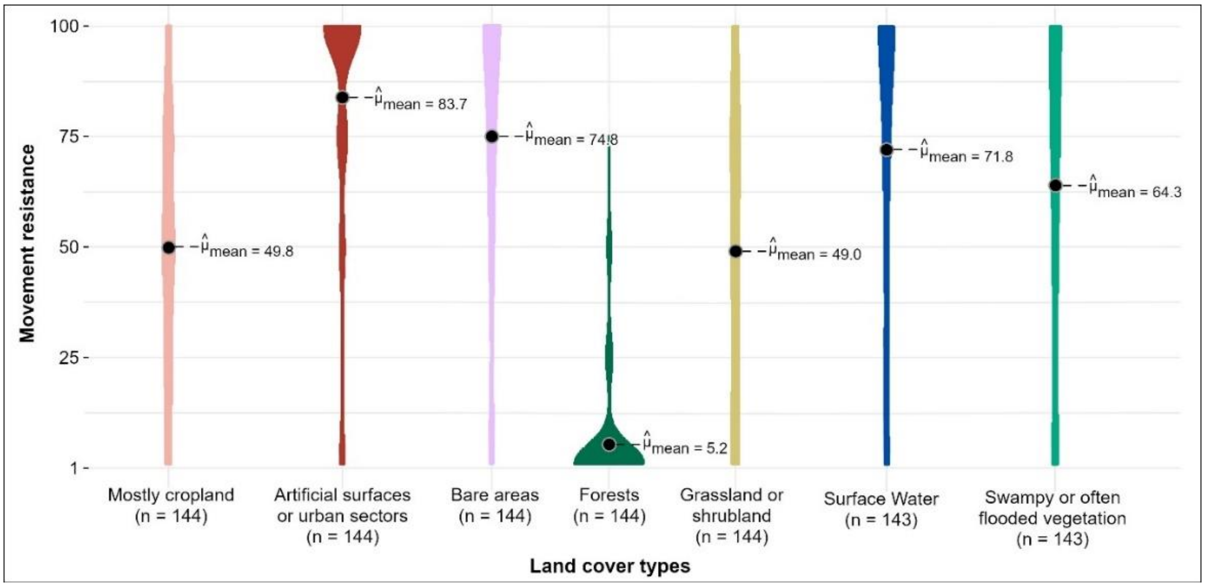


Figure A3. Distribution of expert-rated resistance to movement values for all 26 focal bird species and seven land cover classes considered.
