## Supplementary Material C for "Prioritizing ecological connectivity among protected areas in Colombia using a functional approach for birds"

**Validation**

To validate our corridor models, we used an independent occurrence dataset for each species obtained from eBird, including only records taken after August 2021 and cleaned using the same procedure described in the SDM section. Although occurrences can include non-movement data, they are valuable for connectivity analysis as they provide a quick and inexpensive way to verify model expectations when dispersal movement information is unavailable (Wade et al., 2015). Therefore, for this validation, we predicted that species would be more likely to be recorded in areas with high connectivity values or PAs with sufficient area (Lalechère & Bergès, 2021; McClure et al., 2016). We quantified the proportion of occurrences for each individual species and groups of species located within their respective selected PAs and the highest 20% and 50% corridor values (Ersoy et al., 2019; Mariela et al., 2020; McClure et al., 2016).

At least 54% of the focal species records were located within the top 50% corridor values and PAs identified for each species. The group of birds with small home ranges presented the highest percentage of overlap, while the group with large home ranges showed the lowest.

Table S1. Percentage of focal species occurrences overlapping with Least Cost Corridors (LCCs) and selected Protected Areas (PAs) for the respective species group.

| Species group | Number of species | Number of occurrences | Percentage of records overlapping top 20% corridor values and PAs | Percentage of records overlapping top 50% corridor values and PAs |
| --- | --- | --- | --- | --- |
| Birds with small home ranges | 14 | 4,689 | 48.5% | 73.2% |
| Birds with medium home ranges | 7 | 656 | 42.2% | 69.5% |
| Birds with large home ranges | 5 | 573 | 33.8% | 54.27% |
| All species | 26 | 5,929 | 43.5% | 68.1% |
